# PhenoStream: A Cyberinfrastructure for Automated and AI-Based Crop Trait Extraction from Aerial Imagery

**DOI:** 10.64898/2026.08.26.747008

**Authors:** Sebastian Varela, Jeremy Ruhter, Erik Sacks, Xuying Zheng, Dylan Allen, Anna Hale, Cory Landry, Xianyan Kuang, Blake Long, Ernst Cebert, Yan Zhu, Shatabdi D. Proma, Sehijpreet Kaur, Diego Jarquin, Jesse Morrison, Andrew D.B. Leakey

## Abstract

The integration of digital technologies for high-throughput field phenotyping is critical for accelerating crop improvement in agriculture. However, extracting traits from remote sensing data remains constrained by fragmented workflows, manual intervention, and limited interoperability among existing tools, resulting in delays that hinder timely biological insight and decision-making. To address these challenges, we present PhenoStream (Phenotyping Streaming), a scalable, end-to-end cyberinfrastructure designed to automate the full lifecycle of aerial imagery–based phenotyping, from data acquisition to plot-and genotype-level inference. The framework integrates automated data ingestion from distributed field sites, geospatial processing, and AI-enabled trait extraction within a unified, user-accessible graphical interface. Its modular and extensible architecture supports adaptable trait modeling and seamless integration of new data sources, enabling deployment across diverse crops, environments, and experimental designs. We demonstrate the system across a large multi-location field trial network of bioenergy crops, where it enables high-throughput characterization of spatiotemporal growth dynamics, genotype-by-environment (G×E) interactions, and predictive modeling of key agronomic traits. By significantly reducing processing latency and manual effort, the platform facilitates near-real-time analysis and reproducible workflows. This work establishes a generalizable and scalable pathway for operationalizing very-high-spatial resolution aerial phenotyping in agricultural research. By bridging data acquisition and analytics, the end-to-end cyberinfrastructure provides a foundation for integrating heterogeneous and unstructured data streams—including remote sensing, environmental, and management data—toward data-driven decision making in agriculture.

## INTRODUCTION

Developing high-yielding and resilient bioenergy cultivars tailored to diverse climatic regions and end-use applications, requires the selection of genetic materials optimized and adapted to local conditions (Khaipho-Burch et al., 2023). A fundamental challenge in crop breeding programs is the accurate and timely characterization of complex traits—such as chemical composition, growth rate, and yield—that are shaped by both genetic factors and environmental variability (Baxter et al., 2012) (Edwards et al., 2016), (Clark et al., 2019). Traditional phenotyping methods, which rely on manual collection of morphological and agronomic traits, remain labor-intensive, slow to collect, and inefficient for large-scale, multi-environment trials analysis (Varela et al., 2022b).

Recent advances in high-throughput phenotyping technologies, particularly remote sensing, provide scalable alternatives for monitoring crop performance across space and time (Varela et al., 2021). Aerial imaging platforms, including uncrewed aerial vehicles (UAVs) and satellites, enable frequent, non-destructive monitoring of entire field experiment trials, substantially reducing labor requirements while improving both spatial resolution and temporal revisit frequency (Ranđelović et al., 2023), (Varela et al., 2025). These data streams enable detailed modeling of crop responses to environment variability, enhancing the precision of selection and management decisions (Araus and Cairns, 2014).

Despite these advances, a major bottleneck remains in transforming sensor data into actionable biological insights. Existing workflows are often fragmented, requiring multiple incompatible software tools and substantial manual intervention to process large datasets (Chen and Zhang, 2020), (Wang et al., 2024) (Varela et al., 2022a). Furthermore, the absence of standardized and scalable data-processing pipelines fosters data silos, hindering reproducible analyses and constraining the range and complexity of artificial intelligence (AI) models that can be effectively implemented, tested, and deployed for practical use (Machwitz et al., 2021).

A critical need therefore exists for robust, and automated computational frameworks capable of managing the full phenotyping data lifecycle—from data acquisition and transfer to processing, modeling, and generation of actionable insights (Yang et al., 2020). Integrating high-resolution aerial phenotyping into breeding and agronomic workflows has the potential to enable continuous monitoring, accelerate the identification of genotypes with both local adaptation and broad stability, and support G×E analyses (Bhandari et al., 2023) (Kar et al., 2020). However, realizing this potential requires moving beyond isolated tools and proof-of-concept studies toward interoperable cyberinfrastructure capable of supporting large-scale, multi-location experimentation and timely decision-making. This need is particularly relevant in resource-constrained scenarios, such as in the academic sector, where access to scalable operational phenotyping remains limited.

Addressing these challenges requires the development of integrated systems that minimize human intervention, reduce reliance on proprietary tools, and provide scalable, reproducible, and user-accessible analytical capabilities (Gano et al., 2024). Fully realizing the potential of sensing technologies requires moving beyond proof-of-concept studies and isolated software solutions toward scalable computational infrastructure capable of managing the entire data-to-insights continuum, including automated data ingestion and secure transfer from remote locations, standardized processing, modeling, and dissemination of inference-ready insights. Without such a foundation, even advanced sensing and modeling approaches remain limited in their practical impact.

In this study, we present PhenoStream, an and-to-end cyberinfrastructure designed to automate and accelerate aerial imagery-based phenotyping workflows. The proposed system integrates data transfer, centralized storage, geospatial processing, and AI-enabled trait inference within a unified computational framework. Its modular architecture supports phenotypic traits extraction and facilitates the characterization of spatiotemporal growth dynamics and G×E interactions across diverse experimental conditions.

The system is demonstrated using a multi-location network of field trials in the context of bioenergy crop breeding programs across a latitudinal gradient in the United States. This use case provides a large-scale, heterogeneous testbed for evaluating the cyberinfrastructure’s ability to support distributed data acquisition, processing, and analysis workflows. The proposed solution automates data transfer from remote operations, centralizes data management, and enables scalable analytics through interactive web-based geospatial tools. By addressing traditional data silos and workflow fragmentation, the framework supports high-throughput and reproducible analysis spanning from raw imagery to plot-level insights. Additionally, the proposed solution facilitates downstream analyses, including characterization of G×E interactions and AI-based predictive modeling on critical phenological stages and yield.

This paper is organized as follows: (1) description and implementation of the cyberinfrastructure; (2) overview of data collection and multi-location field network; and (3) downstream analyses enabled by the system, including spatiotemporal dynamics of genetic material across environments, predictive modeling of key traits, and an example use case in the context of plant breeding.

## MATERIALS AND METHODS

### Cyberinfrastructure Description and Implementation

PhenoStream was developed as centralized, automated data management system enabling distributed data ingestion and integrated processing of aerial imagery for plant phenotyping applications.

### LEMP Web Application

The backbone of the system is implemented as a full-stack Linux, Nginx, MySQL, and PHP (LEMP) web application deployed on a server at the University of Illinois, Urbana-Champaign, supporting seamless data transfer from geographically distributed field sites to a centralized processing environment (Fig. 1A-C). This component is responsible for initial data ingestion, validation, storage, and photogrammetric processing.

**Figure 1.**
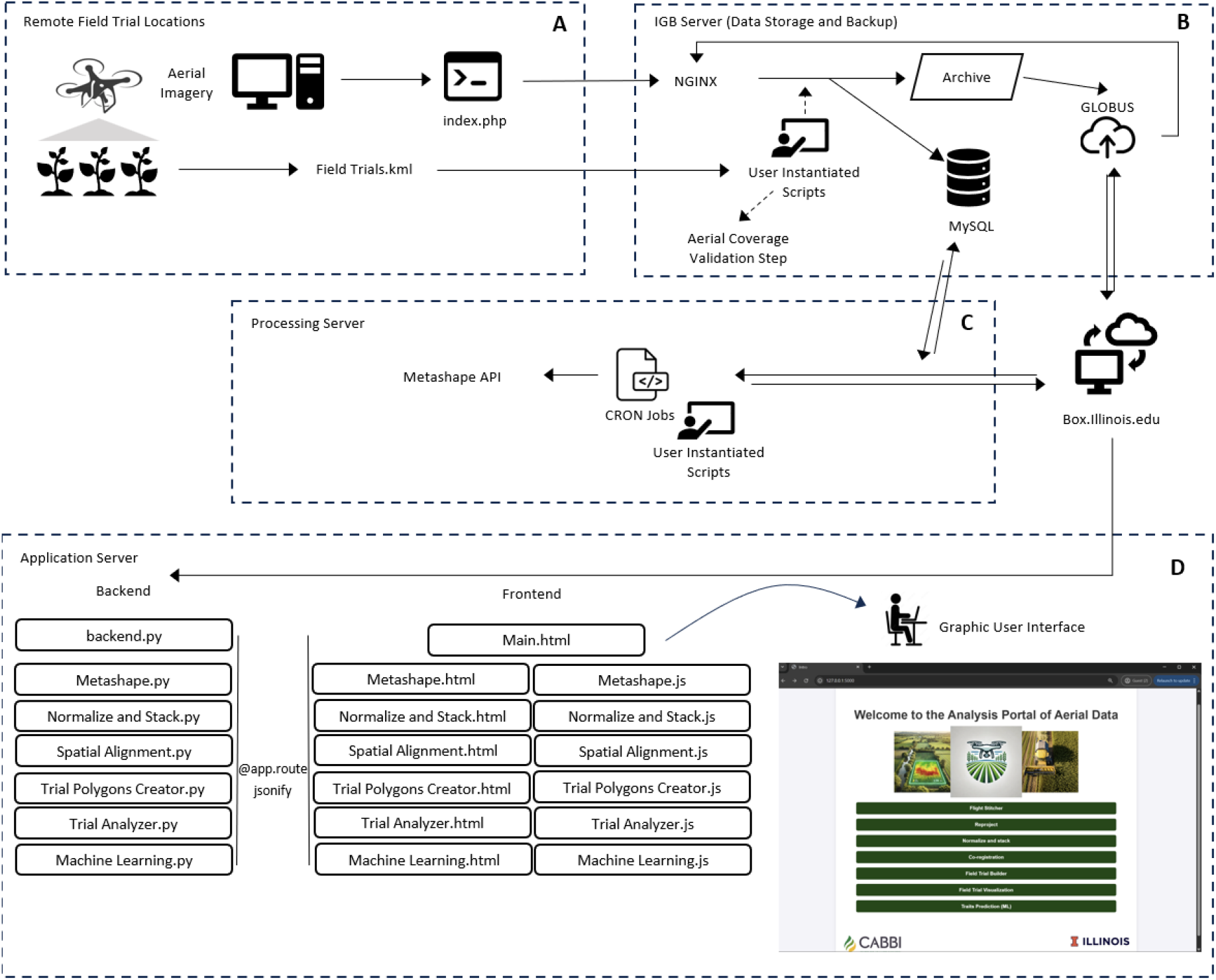
Overview of PhenoStream framework for data ingestion, processing, and analysis. (A-C) Centralized LEMP infrastructure for automated data management and processing: (A) data acquisition and automated upload to the central server via SD card scripting; (B) validation of aerial coverage and multi-tier data management, including local storage, MySQL database integration, cloud backup (Box), and automated data transfer via Globus; and (C) automated image processing pipeline executed through CRON jobs using the Metashape Python API, generating multispectral orthomosaics and CSMs. (D) Modular front–backend analytical environment for interactive geospatial analysis, enabling transformation of field-scale raster outputs into plot-level insights through web-based interfaces that execute backend Python routines.

A PHP-based Dropzone interface facilitates efficient upload of imagery collected at remote location (Fig. 1A). Upon upload, a validation pipeline automatically verifies image metadata against predefined GNSS-referenced field boundaries, associating each dataset with its corresponding experiment and ensuring proper data organization. In parallel, a custom PyCUDA-based routine evaluates spatial coverage using image geopositioning metadata to confirm completeness of field acquisition (Fig. 1A and B). Following validation, images enter a triple-tier backup workflow consisting of (1) a local server archive, (2) a MySQL database, and (3) an external cloud storage (Box), with data transfers managed via Globus (Fig. 1B). The LEMP application also provides a custom PHP search engine for querying and retrieving processed datasets. A scheduled Globus synchronization process continuously monitors incoming data and ensures secure-off-site data integration.

Processing within the LEMP-based infrastructure is fully automated through scheduled CRON jobs, which trigger pipeline execution for newly ingested datasets (Fig. 1C). Image processing leverages the Agisoft Metashape Python API (Agisoft, St. Petersburg, Russia) to perform photogrammetric processing, including image alignment, spectral calibration, dense point cloud generation, and the production of multispectral orthomosaics and crop surface models (CSMs) on a per-field-trial basis. Final outputs are systematically stored in both the database and cloud repository, enabling downstream use.

Outputs resulted from these steps (Figure 1A–C) serve as inputs to a modular front–backend analytical environment designed for interactive visualization and advanced analysis (Fig. 1D), described in the following subsections.

### Interactive Analytical Environment

Complementing the backend data management and processing pipeline (Fig. 1A-C), a separate modular analytical environment was developed to enable interactive visualization and downstream geospatial analysis (Fig. 1D) as part of PhenoStream. This layer operates on prior intermediate outputs (e.g., orthomosaics, CSMs) and provides users with tools for transforming field-scale raster data into plot-level insights.

The system integrates a Python-based backend with a dynamic web frontend, enabling seamless execution of geospatial workflows without requiring direct script interaction. The backend, implemented in Flask, manages routing, logic execution, and server-side processing, while the frontend—built with HTML, CSS, and JavaScript—supports interactive visualization and real-time communication via AJAX and JSON. User inputs trigger backend routines that execute analytical tasks and dynamically update outputs (e.g., map layers and plots) without page reloads.

The application is deployed on a dedicated application server and structured around a central interface that routes to multiple analytical modules (Fig. 2A-G). Each module corresponds to a specific stage in the transformation of raster (i.e., orthomosaics and CSMs) data into structured, analysis-ready datasets. each connected to a corresponding Python script called from *back.py* (Fig. 1D). The architecture enables scalable integration of geospatial processing, feature extraction, and AI-based inference within a unified analytical workflow.

**Figure 2.**
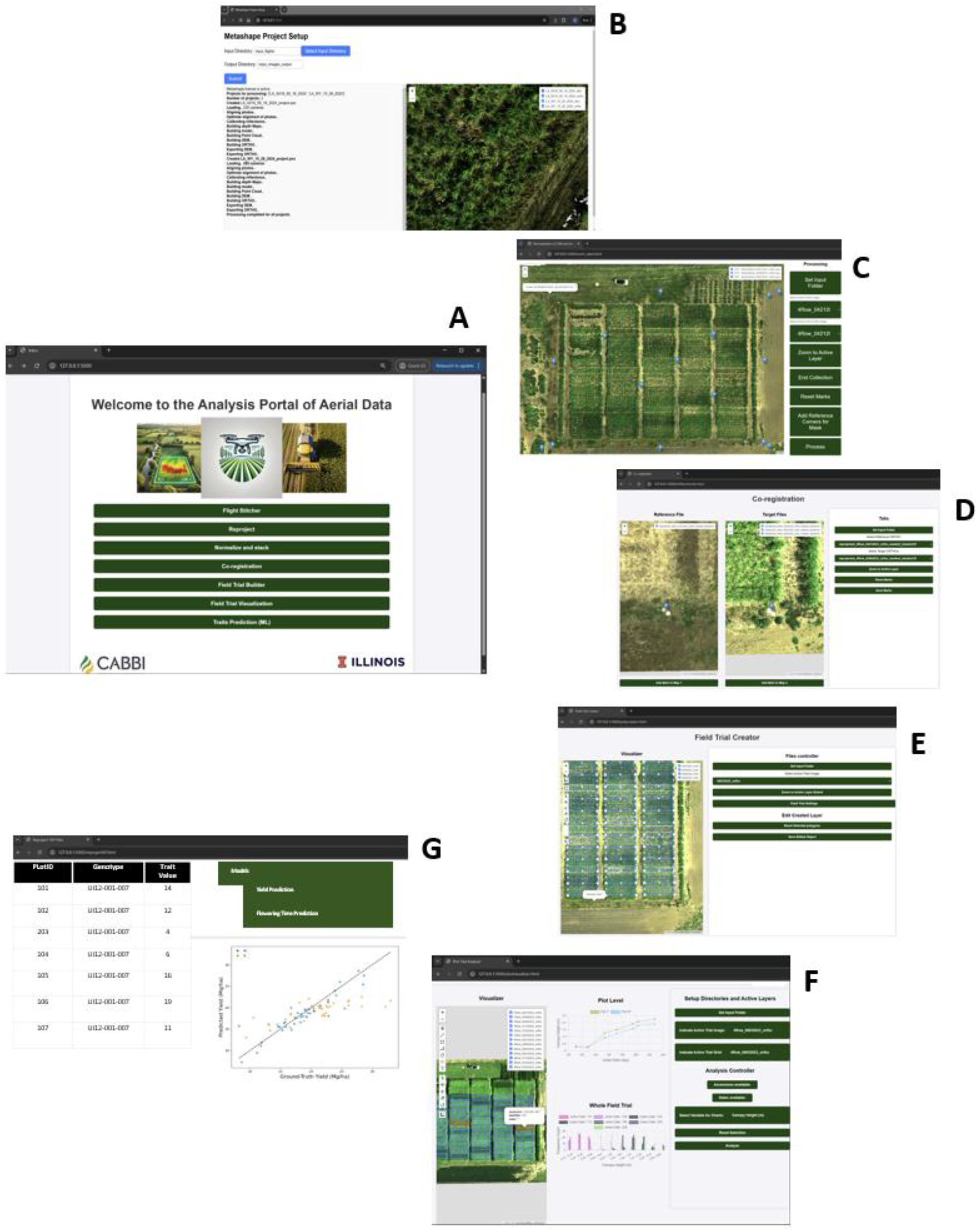
Modular analytical components of the Interactive Analytical Environment for transforming raster data into plot-level insights. (A) Main landing page providing access to each of the analytical modules. (B) Image Processing and Reconstruction module for interactive photogrammetric processing and orthomosaic and CSM generation. (C) Preprocessing and Feature Standardization module, including terrain normalization, masking, and multispectral layer stacking. (D) Spatial Alignment and Temporal Consistency module for ensuring alignment across multiple acquisition dates within the same field trial. (E) Field Trial’s Plots Delineation module for encoding experimental layout design through multi-polygon generation. (F)

A dedicated module provides a graphical user interface (GUI) for batch processing using the Agisoft Metashape Python API (Fig. 2B). Although photogrammetric processing is primarily managed throughout the LEMP application workflow (Fig. 1C), this interface provides a flexible alternative for photogrammetric processing without requiring coding expertise. The module automates image alignment, camera optimization, reflectance calibration, dense point cloud generation, and the creation of orthomosaics and CSMs. It is particularly useful for batch processing UAV flight datasets organized in date-based subfolders, enabling progress monitoring and visualization of outputs through an integrated map interface.

Plot-Level Feature Extraction and Analysis module for rapid exploratory assessment of temporal dynamics at plot or genotype levels. (G) AI Analyzer module for predictive modeling of traits such as flowering dynamics and yield.

### Preprocessing and feature standardization

This module prepares raster datasets for downstream analysis by ensuring spatial and structural consistency (Fig. 2C). It identifies corresponding orthomosaic and CSM date pairs, performs terrain normalization using a k-nearest neighbors interpolation of ground points, and applies masking and resampling to standardize spatial extent and resolution. Spectral and structural layers are then stacked into unified raster objects, enabling consistent further analysis.

### Spatial alignment and temporal consistency

To support time-series analysis, a spatial alignment module was developed to correct residual positional inconsistencies across acquisition dates (Fig. 2D). Despite Real Time Kinematics (RTK)-enabled acquisition, minor spatial offsets between dates can affect temporal consistency and represent a significant technical bottleneck for temporal analyses of consecutive flights, particularly when using non-RTK UAV systems. The module groups raster datasets by trial and acquisition date, allowing users to define a reference layer to which subsequent datasets are aligned through manual or semi-automated adjustment options (Fig. 2D). The resulting aligned datasets ensure consistent plot-level temporal feature extraction, which is critical for downstream analyses.

### Field trial’s plots delineation

This module converts field-scale raster data into structured experimental units by generating multi-polygon representations of field trial layouts (Fig. 2E). Using an interactive map interface, users define plot geometry, orientation, and experimental design parameters (e.g., row/column structure, block configuration). The resulting spatial objects encode the experimental design and enable consistent spatial linkage between the raster data and the plot-level observations done on the ground. Prior to finalizing and saving the trial layout object as JSON file, an editable intermediate step allows manual adjustment of the multi-polygons relative to the underlying RGB orthomosaic to ensure accurate spatial alignment.

### Plot-level feature extraction and analysis

This component enables an interactive exploratory analysis of temporal dynamics at the plot and genotype levels (Fig. 2F). Users can visualize plot and genotype temporal trajectories and statistical summaries of derived traits, such as canopy height and vegetation indices. Two Interactive figures support dynamic filtering and comparison across genotypes and time points, enabling rapid identification of patterns, trends, and outliers.

### AI analyzer

A dedicated module supports AI–based trait inference using plot-level datasets (Fig. 2G). Users can input imagery and associated ground-truth data to perform predictive tasks such as yield estimation or determining flowering dynamics. The GUI enables model selection and visualization of outputs, including predicted versus observed relationships or temporal progression patterns. This component integrates advanced analytical methods into the workflow while maintaining accessibility for non-coding users.

### Multi-location Field Network and Data Acquisition

To implement PhenoStream into real-world conditions, the system was deployed across a multi-location network of field trials spanning four locations in the central and southeastern United States (Fig. 3). The network comprises 31 field trials monitored between 2020 and 2024, covering a latitudinal gradient of approximated 11 degrees and encompassing diverse environmental conditions.

**Figure 3.**
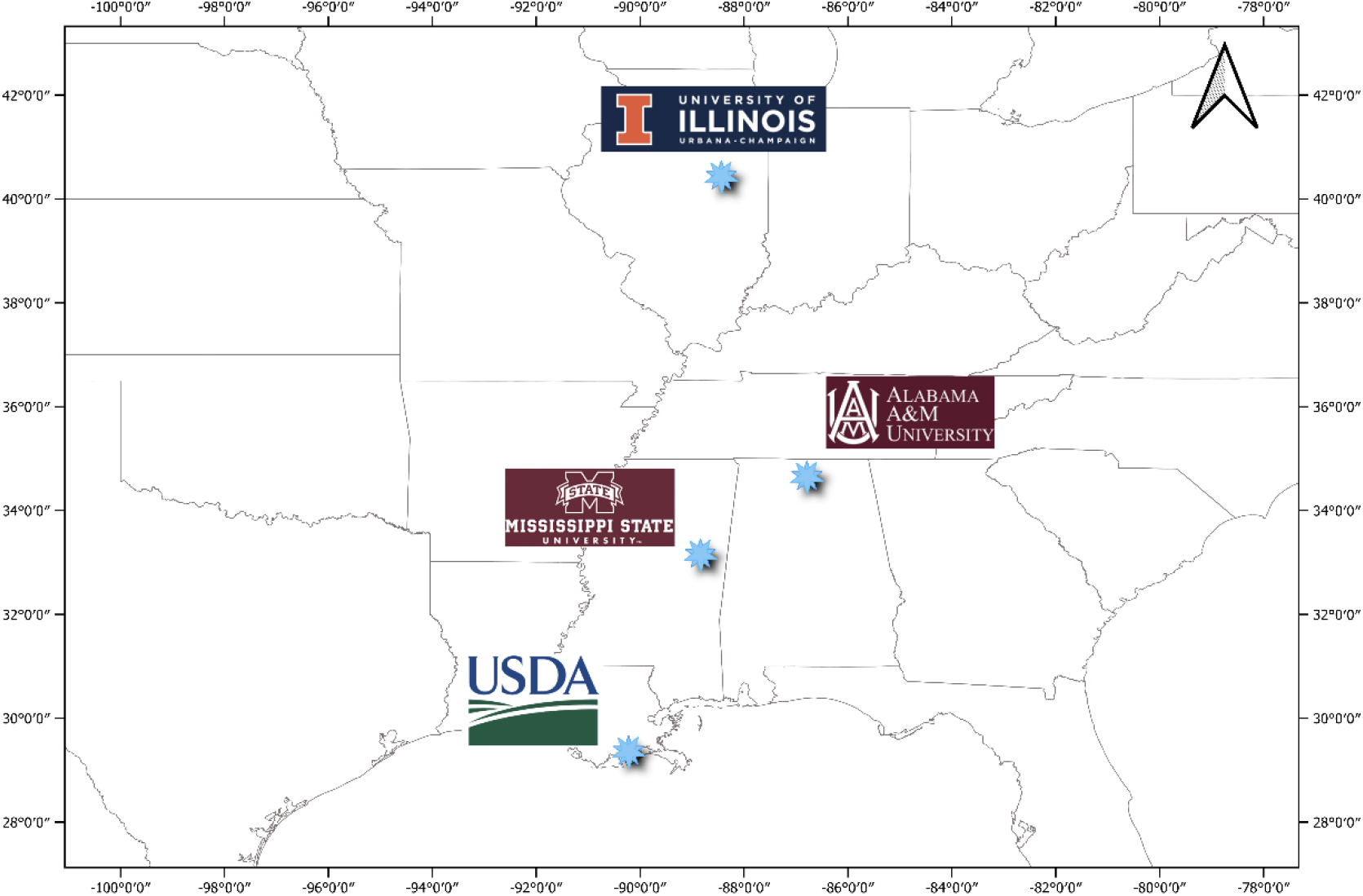
Geographic distribution of the network sites across the central and southeastern United States. Sites include: the Energy Farm Research Station at the University of Illinois Urbana-Champaign (Urbana, Illinois, 40°03’56”N, 88°12’30”W), the Winfred-Thomas Agricultural Research Station at Alabama A&M University (Hazel Green, Alabama, 34°54’00”N, 86°33’41”W), the MAFES Bearden Dairy Research Center at Mississippi State University (Starkville, Mississippi, 33°24’09”N, 88°44’33”W), and the USDA Research Station (Houma, Louisiana, 29°38’05”N, 90°50’27”W).

The field trials at each site span multiple bioenergy crops species, including *Miscanthus* spp., interspecific hybrids, and energycane, enabling evaluation of genotype performance across contrasting agroecological zones. Northern locations are primarily focused on *Miscanthus* species, while southern sites incorporate both *Miscanthus* and energycane, supporting the assessment of crop adaptation from temperate to subtropical environments. This configuration provides a large-scale, heterogeneous dataset suitable for analyzing G×E interactions and testing the scalability of the proposed of the computational infrastructure. Detailed descriptions of sites and trials are provided in the Supplementary Material, with a summary in Table S1.

A Phantom 4 multispectral RTK platform (DJI, Shenzhen, China) was used across all sites to ensure consistent, high-resolution data acquisition. The system integrates RTK positioning with five spectral bands: blue (450 nm), green (560 nm), red (650 nm), red edge (730 nm), and near-infrared (840 nm), each with a bandwidth of ±16 nm, as specified by DJI. Flights were conducted approximately biweekly during the growing season, following a standardized protocol for flight planning, acquisition settings, and data handling to ensure reproducibility and cross-site consistency (Fig. 4). Detailed information on platform selection, data collection, and operational implementation is provided in the Supplementary Material.

**Figure 4.**
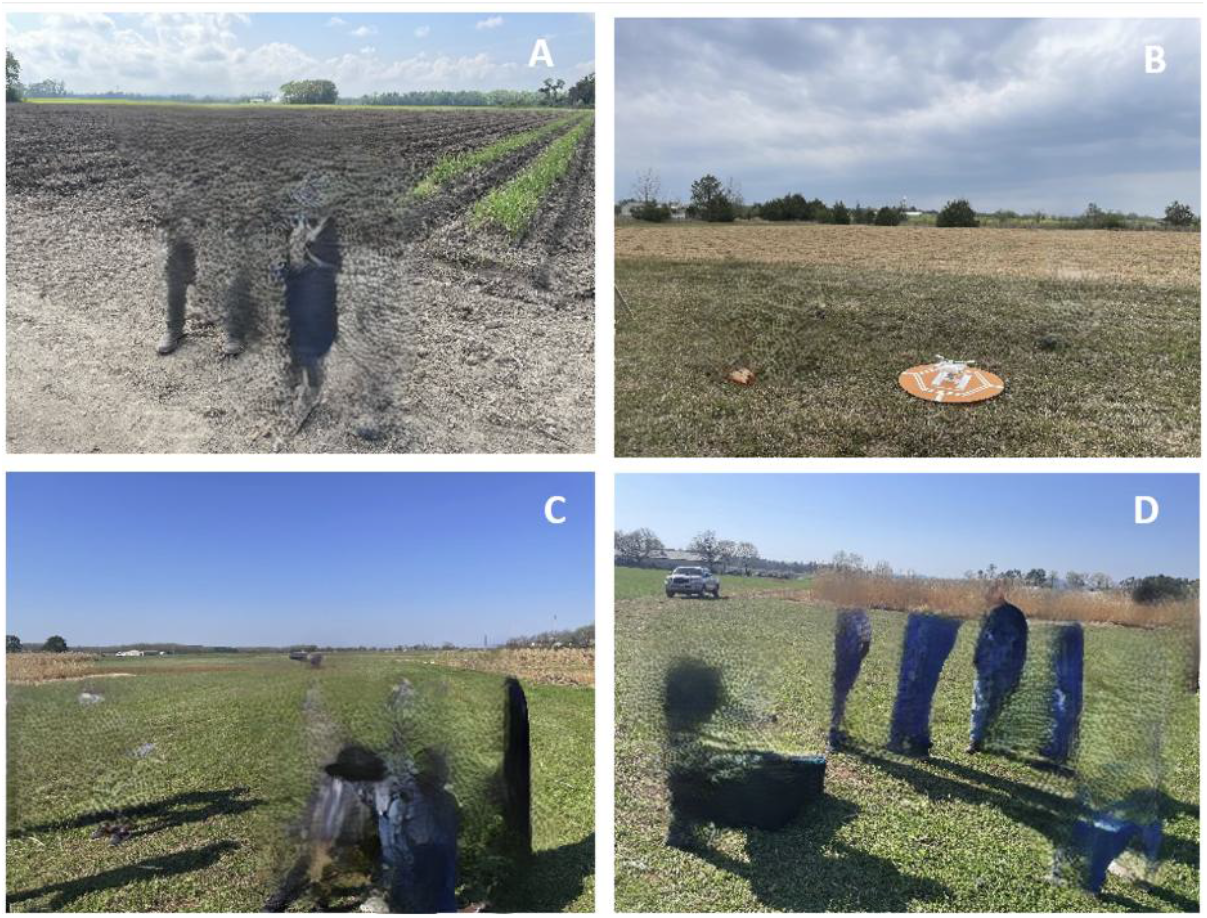
Training and coordination activities conducted to standardize aerial data collection across network sites (identifying information of people was removed as requested by bioRxiv). (A) Illinois team discussing data acquisition protocols with the Louisiana team at the USDA Research Station in Houma, LA. (B) Example of flight-planning demonstration conducted by the Illinois team at the Mississippi site. (C–D) Interaction between the Illinois and Alabama teams at the Winfred-Thomas Agricultural Research Station in Huntsville, AL, in March 2023.

A total of 560 aerial flights were conducted between 2020 and 2024 across the four locations (Fig. 5), generating 4.8 TB of raw imagery. Data collection was initially limited to Illinois (IL), with 39, 23, and 18 flights conducted in 2020, 2021, and 2022, respectively. The network expanded in 2023 to include Alabama (AL) (79 flights), Mississippi (MS) (61), Louisiana (LA) (26), and IL (20), and further increased in 2024 with 146, 74, 43, and 31 flights at these locations, respectively. This progression reflects the transition from a single-location deployment to a distributed multi-location network, substantially increasing spatial and temporal coverage. Flights were aligned with key crop phenological stages, spanning early vegetative growth through flowering, with day-of-year coverage varying by location and crop type. Data were processed through PhenoStream. Raw imagery was first transformed into high positional accuracy orthomosaics and CSMs. These outputs were then aggregated into a structured, longitudinal dataset capturing multi-year, multi-location crop dynamics at high temporal resolution. The dataset was subsequently integrated with AI-based trait inference for downstream applications.

**Figure 5.**
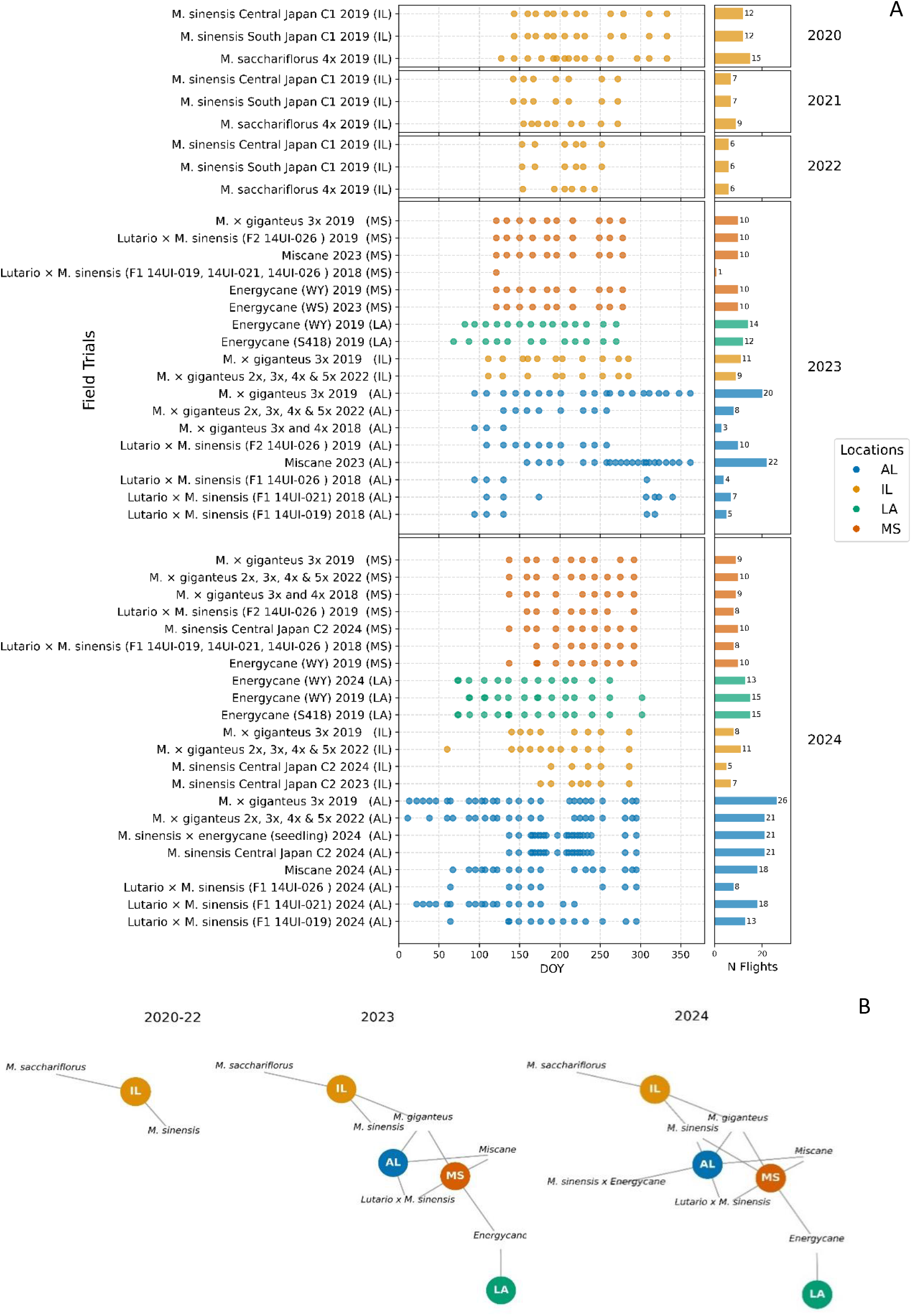
(A) Summary of data collection across network locations, including field trial names (left) and the temporal distribution of flights (center) by year (top to bottom) across locations (colors). (B) Distribution of flights across locations by species and year (left to right).

## RESULTS

### Downstream Entry-Level Growth Dynamics Analysis

Species-level temporal dynamics coherently reflected the latitudinal gradient across the network, with earlier green-up and steeper growth trajectories observed in southern locations compared to northern sites (Fig. 1S). These patterns were consistent across years and species, establishing the environmental baseline within which genotype-level differences were evaluated. A more detailed species-level temporal dynamics description is provided in Supplementary Materials.

To evaluate the sensitivity of PhenoStream at a fine scale, temporal dynamics derived from the pipeline (Fig. 1) were examined at the entry level in two field trials from different species— Miscanthus × giganteus 3x 2019 and Energycane (WY) 2019 trials —across network locations. Analyses focused on two entries, Miscanthus × giganteus 3x 2019 UI12-001-007 and Energycane (WY) 2019 Ho14-9213, using three complementary approaches: (i) visualization of temporal dynamics by location and year with canopy height (CH) and normalized difference vegetation index (NDVI), (ii) quantification of temporal dissimilarity within and between environments via pairwise distance analysis (Gower, 1971) based on CH and NDVI, and (iii) characterization of phenotypic plasticity across environments (i.e., G × E interactions) using reaction norms (Schlichting and Pigliucci, 1998) with CH.

Pairwise distance (Virtanen et al., 2020) was calculated as the Euclidean distance between plots’ CH and NDVI values, where each plot represents a replicate of an entry in a given trial and location. Distances were grouped into within-location (plots of the same entry within a location) and between-location (across locations) categories to assess how well phenotypes clustered by environment (Location × Year). Reaction norms were constructed as line plots of mean CH values per genotype across environments: parallel lines indicate marginal G × E (consistent performance across locations), whereas crossing lines indicate strong G × E (environment-dependent performance).

The temporal dynamics of UI12-001-007 (Fig. 6A-B) broadly followed the patterns observed for the species across locations (Fig. S1E-F). In AL, green-up occurred earliest and steepest, plateauing earlier than in other sites (Fig. 6A-B). In IL, green-up was delayed but eventually converged with MS (DOY 250 for CH, DOY 150 for NDVI). MS green-up began slightly earlier than IL but progressed more slowly, resulting in consistently lower values throughout the season. Notably, the entry’s CH and NDVI slopes and absolute values in AL and IL (Fig. 6A-B) appear visually higher than the overall entry means (Fig. 6A–B). Across years, UI12-001-007 exhibited interannual variability (Fig. 6A–B). In AL, CH and NDVI were slightly higher in 2023 than in 2024, likely reflecting environmental or management effects (e.g., weather, harvest timing). In IL, CH increased more rapidly in 2024, whereas in MS both traits showed persistently slow growth.

**Figure 6.**
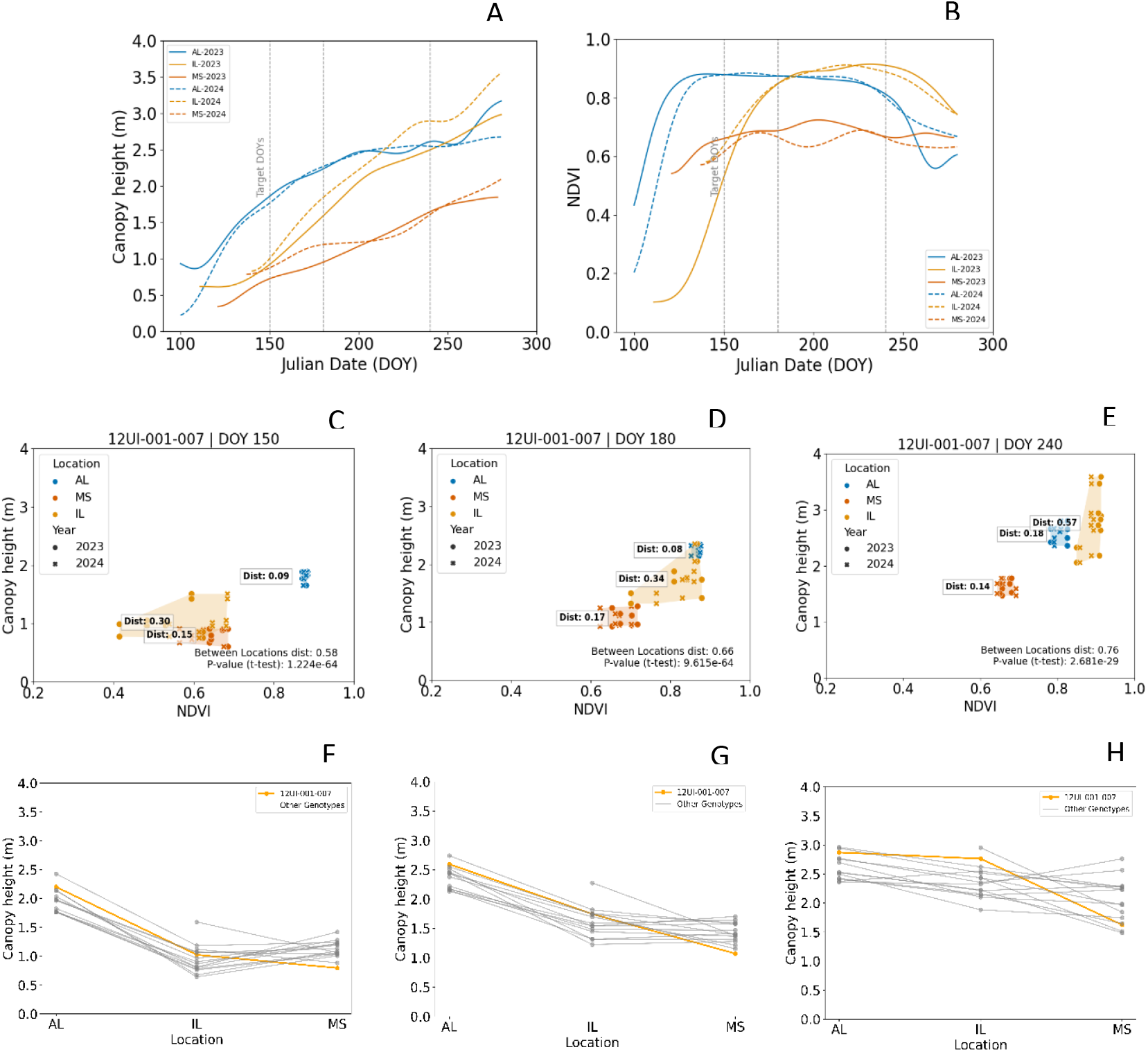
Integrated analyses of Miscanthus × giganteus 3x 2019 12UI-001-007 entry, planted in 2019 at locations in AL, IL, and MS, with data collected in 2023 and 2024. (A–B) Mean temporal trajectories of 12UI-001-007 for CH and NDVI. (C–E) Pairwise distance analyses based on CH and NDVI at DOYs 150, 180, and 240. (F–H) Reaction norm analyses of entry 12UI-001-007 (orange line) compared with other entries (grey lines) at DOYs 150, 180, and 240.

Pairwise distance analyses confirmed smaller dissimilarity within locations than between them (Fig. 6C-E). At DOY 150, within-location distances were 0.09 (AL), 0.30 (IL), and 0.15 (MS), compared with 0.58 between locations. Later in the season (Fig. 6D-E), within-location distances ranged from 0.08 to 0.57, whereas between-location distances were consistently higher (0.66–0.76).

Reaction norms further contextualized UI12-001-007 relative to other Miscanthus × giganteus 3x 2019 entries in the 2019 trial (Fig. 6F-H). As expected, most entries (grey lines) aligned with general CH and NDVI dynamics for the hybrid (Fig. S1E-F), AL entries showed early and rapid growth, IL had delayed but steeper catch-up later in the season, and MS entries showed intermediate onset but weaker growth overall. In contrast, UI12-001-007 (orange line) diverged from this general pattern, ranking consistently high in AL and IL (Fig. 6F-G), but performing relatively low in MS (Fig. 6H).

The entry Ho14-9213 exhibited temporal dynamics consistent with general energycane trends across the two trial locations (MS and LA; Fig. S1G-H). In LA, green-up occurred earlier and with higher slope rate, reflected in consistently stronger CH and NDVI values throughout the season (Fig. 7A-B). In MS, green-up was delayed by several weeks and rose more gradually, though by DOY 280 CH values approached those observed in LA. Interannual variability was evident: in LA, both CH and NDVI were higher in 2024 than 2023, whereas in MS the opposite was observed, with 2023 showing stronger early growth and higher values across the season.

**Figure 7.**
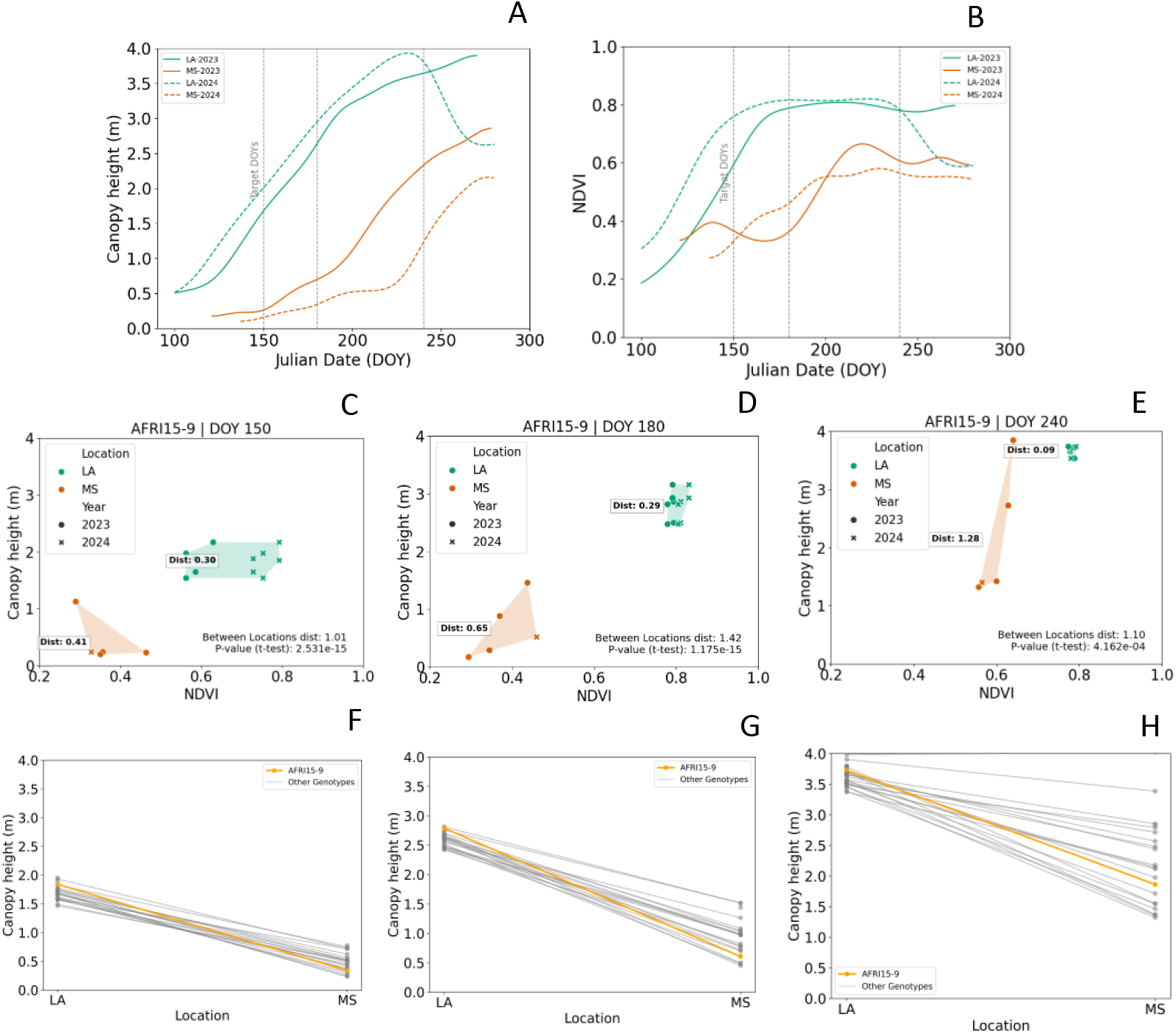
Integrated analyses of Energycane (WY) 2019 Ho14-9213 entry, planted in 2019 at locations in AL, IL, and MS, with data collected in 2023 and 2024. (A–B) Mean temporal trajectories of Ho14-9213 for CH and NDVI. (C–E) Pairwise distance analyses based on CH and NDVI at DOYs 150, 180, and 240. (F–H) Reaction norm analyses of entry Ho14-9213 (orange line) compared with other entries (grey lines) at DOYs 150, 180, and 240.

Pairwise distance analysis confirmed that entry dynamics were more similar within locations than across them (Fig. 7C-E). At DOY 150, distances were 0.30 (LA) and 0.41 (MS), compared to 1.01 between locations. At DOY 180, within-location distances were 0.29 (LA) and 0.65 (MS), versus 1.42 between locations. By DOY 240, values still low 0.09 (LA) and increased to 1.28 for MS, while the between-location distance was 1.10. This indicated a larger distance in MS than between locations; however, this was mainly driven by one of the repetitions of the entry which exhibited an extreme large value in MS which inflated the within MS distance value.

Reaction norms further contextualized Ho14-9213 relative to other entries in the Energycane (WY) 2019 trial (Fig. 7F-H). Overall, entry trajectories (grey lines) aligned with species-level dynamics (Fig. S1G-H), with LA consistently outperforming MS across the season. MS also displayed greater variability from mid-season onward (DOY 180; Fig. 7G-H). Within this context, Ho14-9213 (orange line) followed the general trend of higher performance in LA than MS, ranking among tops entries in LA, while only at mid-low range in MS location (Fig. 7H).

### Downstream AI-based Traits Inference

Trait inference based on pretrained algorithms can be implemented to generate plot-level predictions for traits such as yield and flowering progress.

An interactive analyzer enables yield inference (Fig. 8) and is supported by a Vision Transformer (ViT)-like architecture (Dosovitskiy et al., 2021). The model operates on time-point UAV imagery (Fig. 8A) in combination with user-provided ground-truth data. To reduce dimensionality while preserving fine-scale spatial information, each image is partitioned into non-overlapping patches that are flattened and projected into a fixed-dimensional embedding space (Fig. 8B). Learned positional embeddings encode the spatial arrangement of patches, enabling the model to capture canopy structure and spatial relationships within each image.

**Figure 8.**
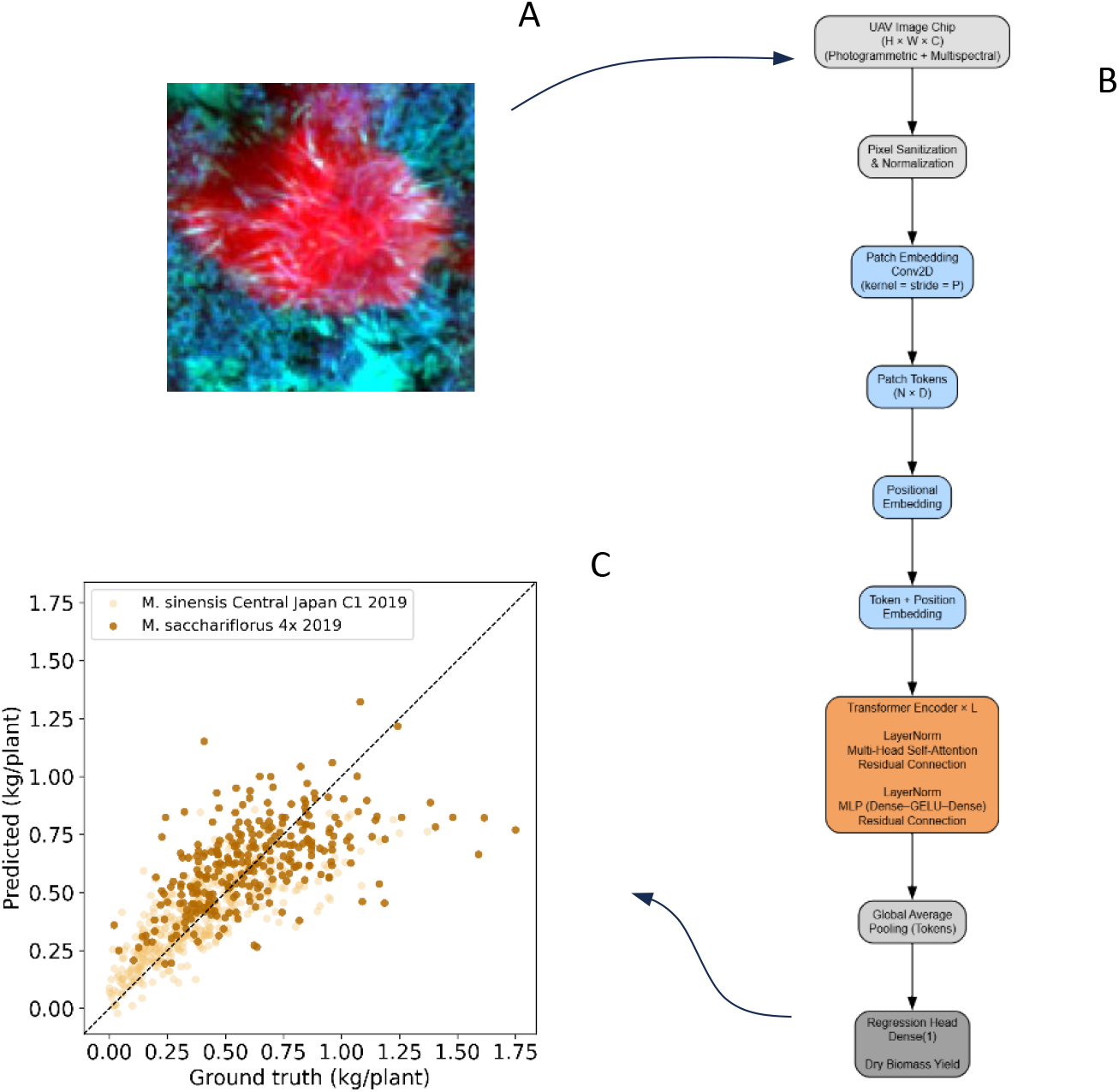
Workflow for AI-based yield prediction inference: (A) The user provides input imagery and ground-truth data; (B) the imagery is then normalized, transformed, and processed by the Vision Transformer (ViT) model; and (C) yield predictions for Miscanthus sacchariflorus 4x 2019 and Miscanthus sinensis Central Japan C1 2019 are subsequently generated and visualized.

The resulting sequence of patch embeddings is processed through a stack of transformer encoder blocks. Within each block, multi-head self-attention models long-range spatial dependencies across the image, allowing the network to jointly represent canopy architecture and localized spectral reflectance patterns. Residual connections and layer normalization promote stable optimization, while a feed-forward multilayer perceptron (MLP) enhances representational capacity. The final patch-level representations are aggregated via global average pooling to produce a compact image-level embedding. This representation is passed to an MLP-based regression head with GELU activation and dropout regularization, followed by a linear output layer that produces trait predictions (Fig. 8C).

The architecture was trained using 9,767 samples with corresponding ground-truth yield measurements from *Miscanthus sacchariflorus* 4× 2019, *Miscanthus sinensis* Central Japan C1 2019, and *Miscanthus sinensis* South Japan C1 2019 populations collected in Illinois during the 2020 and 2021 growing seasons. This training exposed the model to diverse canopy structures, spectral reflectance patterns, and yield across environments and genotypes. Such initialization is expected to facilitate transfer learning by enabling the extraction of generalizable spatial plant features during fine-tuning for downstream prediction tasks. Model performance evaluated on a held-out 30% subset of the 2020 data achieved an R² of 0.55 and a Root Mean Square Error (RMSE) of 187.3.

The second interactive tool enables monitoring of flowering progression backed by a pretrained SimCLR (Simple Framework for Contrastive Learning of Visual Representations) algorithm (Chen et al., 2020) integrated with a binary classifier for inflorescence detection. The tool processes user-provided image chips (Fig. 9A), and ground-truth labels are not strictly required. RGB images are passed through a ResNet50-based encoder with a projection head, producing fixed-dimensional embeddings for each image. The encoder is pretrained using the contrastive SimCLR framework (Fig. 9B). During pretraining, augmented pairs of each image are generated through random flips, rotations, zooms, and contrast adjustments to learn robust representations.

**Figure 9.**
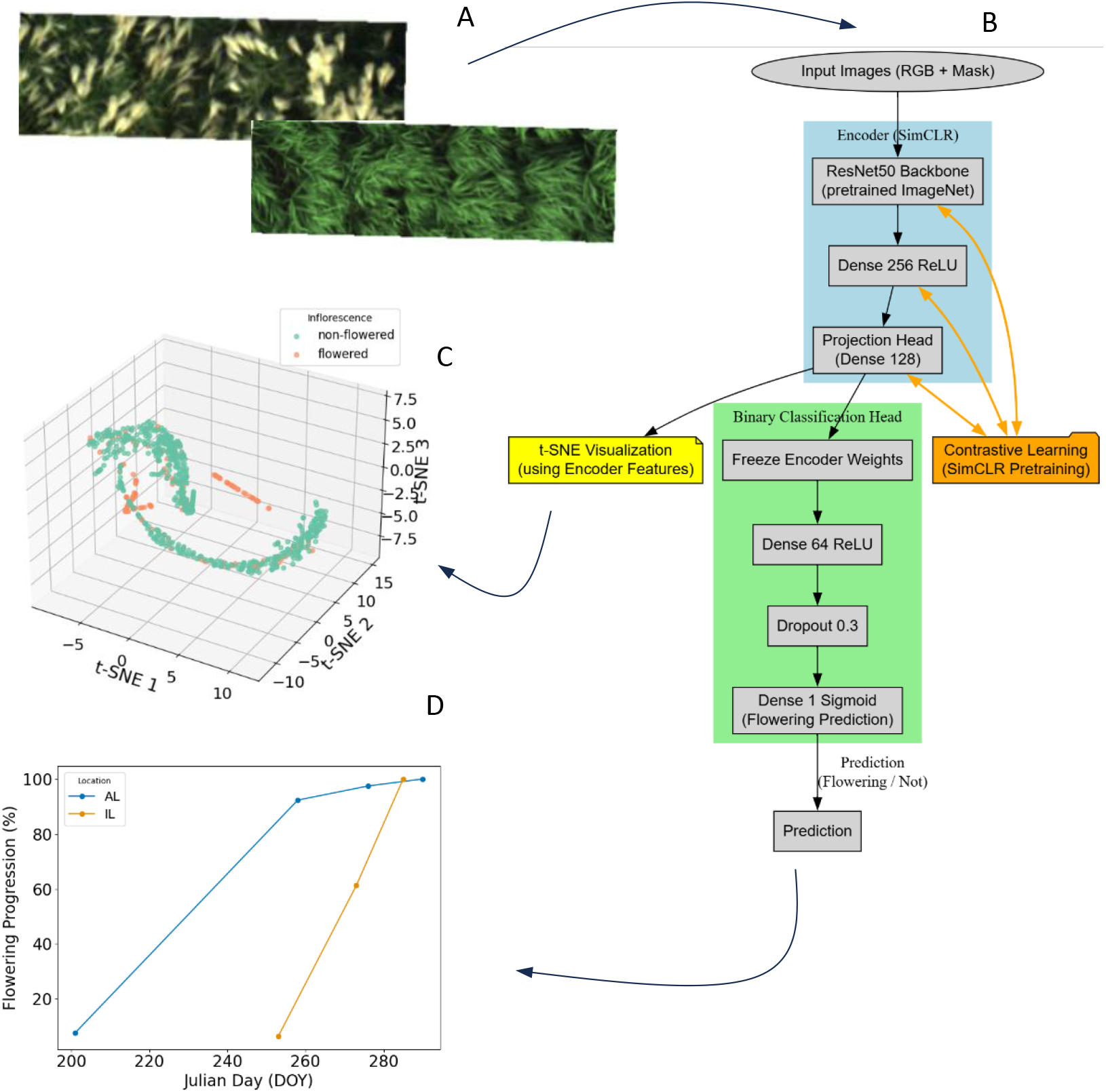
Description of the AI-based workflow for determining flowering dynamics for same entry at two contrasting locations of the regional network, Illinois and Alabama. (A) User provides input RGB imagery with the corresponding date of data collection, and location (environment). (B) Input imagery is normalized, transformed, and passed to the integrated SimCLR and binary classifier model. (C) t-SNE analysis allowing visualization of clusters and separability of image variants by class (i.e., non-flowered versus flowered plants). (D) Flowering progression for Miscanthus × giganteus 3x 2019 at Alabama and Illinois sites.

To explore the structure of the learned embeddings, t-distributed Stochastic Neighbor Embedding (t-SNE) (Maaten and Hinton, 2008) is applied. t-SNE maps high-dimensional embeddings into three dimensions while preserving local similarities, allowing visualization of clusters and separability of image variants by class (i.e., non-flowered versus flowered plants) (Fig. 9C). The 3D t-SNE projections provide insight into how the SimCLR encoder discriminates between image representations.

After pretraining, the encoder is frozen, and a supervised binary classifier is trained on its embeddings to detect the presence of inflorescences. A proof-of-concept implementation was conducted to demonstrate the ability of the framework to detect flowering across a latitudinal gradient. The dataset consisted of 434 images collected in 2023 from *Miscanthus × giganteus* 3× 2019 trials planted in Illinois and Alabama. This implementation demonstrates that accurate flowering detection can be achieved with minimal reliance on labeled data, while maintaining strong classification performance (accuracy = 0.88, precision = 0.93, recall = 0.73, F1-score = 0.78). Once flowering status is predicted, the associated acquisition dates and locations are automatically leveraged to generate a visual representation of flowering progression over time (Fig. 9D).

### Downstream Analysis: A Case Study in Plant Breeding Application

The practical use and modular integration of PhenoStream was evaluated using datasets from Miscanthus sacchariflorus 4x 2019, Miscanthus sinensis Central Japan C1 2019, and Miscanthus sinensis South Japan C1 2019, collected in Illinois during the 2020 and 2021 growing seasons. Dry biomass yield was recorded at the plant level. Populations were analyzed separately. The primary objective was to predict dry biomass yield using UAV-derived features from the cyberinfrastructure pipeline as proxies, under prediction scenarios commonly encountered in plant breeding programs.

Data from multiple UAV flights were integrated using means across flight dates at the plot level for six features (one photogrammetric and five multispectral bands). This strategy enabled the direct integration of information derived from different numbers of flights also collected at different time points across years.

- Single-year main effects model (M1):

This model was used for prediction scenarios restricted to a single year. Dry biomass yield for the *i*th genotype was modeled as:

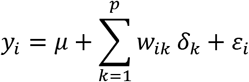

where *μ* is the overall mean, *w_ik_*represents the mean UAV-derived feature *k* for genotype *i*, and *δ_k_*is the corresponding feature effect. The residual error is denoted by *ε_i_*. Assumptions include *δ_k_* ∼ *N*(0, σ^2^_δ_) and *ε_i_* ∼ *N*(0, σ^2^_ε_), with *p* = 6 features.

The phenomic effect for genotype *i* is defined as:

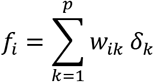

The vector of phenomic effects *f* = {*f_i_*} follows a multivariate normal distribution *f* ∼ *N*(0, Ωσ^2^_*f*_), where **Ω** is the UAV-derived phenomic relationship matrix capturing similarities among genotypes.

- Multi-year main effects model (M2):

To enable predictions across multiple years, the model was extended to include year effects:

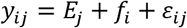

where *y_ij_* denotes the dry biomass yield of genotype *i* observed in year *j*, *E_j_* is the random effect of year *j*, and *ε_ij_* is the corresponding residual error. Year effects follow ***E*** = {*E_j_*} ∼ *N*(**0,Z_*E*_Z_′*E*_σ^2^_*E*_**), where ***Z****_E_* is the incidence matrix linking observations to years.

- Multi-year interaction model (M3):

An additional extension incorporated genotype-by-year interactions:

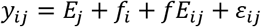

where *fE_ij_* represents the interaction between genotype *i* and year *j*. The interaction term follows ***fE*** = {*fE_ij_*} ∼ *N*(**0**,**Ω**°***Z_E_Z^′^_E_σ^2^_*fE*_***) (Jarquín et al., 2014) and ∘ denotes the Hadamard (element-wise) product.

### Cross-validation schemes

Three cross-validation schemes were implemented to reflect different prediction problems encountered in breeding programs:

#### Within-year prediction

Each year was analyzed independently. A five-fold cross-validation was performed by randomly partitioning the dataset into five non-overlapping subsets. Each subset was used once for validation while the remaining four subsets were used for model training. Predictive ability was assessed as the correlation between observed and predicted values.

#### Forward prediction

Phenotypic and UAV-derived features from 2020 were used for model calibration, and dry biomass yield was predicted for 2021 using UAV-derived features collected in 2021.

#### Across-years prediction

This scheme aimed to predict genotypes not observed in both years while allowing information sharing across years. Genotypes were randomly assigned to five folds. One-fold was used for validation and the remaining four for model training. The procedure was repeated until predictions were obtained for all folds.

The predictive performance of the different models and cross-validation schemes is shown in Figure 10. Predictive ability was evaluated on a yearly basis as the correlation between observed and predicted values. The left panel summarizes results from the within-year and forward prediction scenarios using the single-year main effects model (M1). Within-year predictive ability ranged from 0.73 to 0.82 in 2020 and from 0.64 to 0.68 in 2021 across the three *Miscanthus* populations. Forward prediction exhibited a predictive ability ranging from 0.39 to 0.68. The right panel presents results from the across-years prediction scenario using the multi-year main effects model (M2) and the multi-year interaction model (M3). Across populations and years, the interaction model consistently achieved the highest predictive ability, with values ranging from 0.72 to 0.816 in 2020 and from 0.65 to 0.67 in 2021.

**Figure 10.**
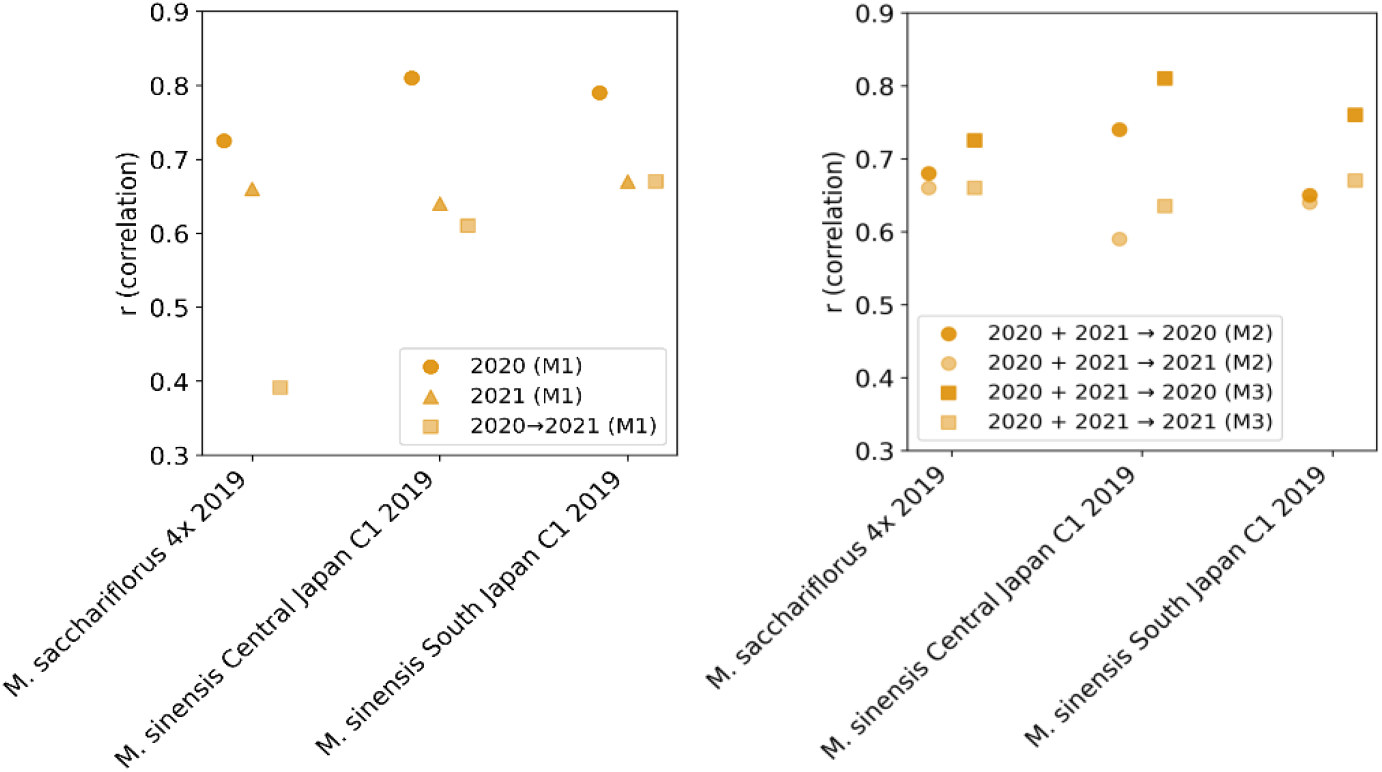
Predictive performance for Miscanthus sacchariflorus 4x 2019, Miscanthus sinensis Central Japan C1 2019, and Miscanthus sinensis South Japan C1 2019 dry biomass yield across three cross-validation schemes (within-year, forward, and across-years). The left panel shows within-year predictions for the 2020 and 2021 growing seasons, as well as forward prediction using 2020 data to predict the 2021 season, all implemented using the single-year main effects model (M1). The right panel presents results from the across-years prediction scenario comparing the multi-year main effects model (M2) and the multi-year interaction model (M3).

## DISCUSSION

The accurate and timely phenotyping of complex agronomic traits across diverse environments is a fundamental bottleneck. This study successfully developed and implemented a cyberinfrastructure that integrates remote sensing, server computing, and AI to minimize data latency and transform raw aerial imagery into rapid and actionable insights across a multi-state regional level. Our framework directly addresses critical challenges of data silos, atomized software solutions, manual processing bottlenecks, and the lack of scalable analytical tools that is limiting the application of high-throughput phenotyping in large-scale, public breeding programs.

The key achievement of this work is the establishment of a robust, end-to-end automated pipeline that significantly reduces the time from data acquisition to biological insight. By moving beyond a collection of disparate software tools to a centralized system with automated uploads, validation, processing, and backup, we have created a reproducible and scalable framework for regional phenotyping. This infrastructure allowed us to process thousands of flights efficiently, a task that would be prohibitively labor-intensive and time-consuming using traditional GUI software or sparse open-source libraries available. The development of a custom geospatial web application with a user-friendly GUI is particularly noteworthy, as it democratizes access to advanced analytical capabilities for breeding teams removing technical and financial barriers (González et al., 2025), enabling them to conduct complex spatiotemporal analyses without requiring expertise in coding or command-line tools. This directly tackles the reproducibility and scalability issues of current approaches.

The spatiotemporal data extracted through this pipeline revealed consistent and quantifiable growth dynamics for key bioenergy crops across a large environmental gradient. The general patterns observed in the features generated with the end-to-end pipeline (e.g., CH and NDVI) align with known phenological behavior for C4 grasses like *Miscanthus*, miscane, and energycane (Ouattara et al., 2022) (Yang et al., 2018). Furthermore, our system captured these patterns with unprecedented spatial and temporal resolution consistency, allowing for fine-scale comparisons.

The clear differentiation of location and year effects, as demonstrated for *Miscanthus × giganteus* and energycane, underscores the powerful influence of environment on phenotypic expression efficiently captured by the downstream analysis of the proposed cyberinfrastructure. For instance, the earlier green-up and steeper growth slope in AL compared to IL and MS for *Miscanthus × giganteus* can be attributed to its more southern, warmer climate. Conversely, the consistently lower CH and NDVI in MS for the same species suggests sub-optimal adaptation or management practices for that specific environment, a hypothesis that requires further investigation. Similarly, energycane’s superior performance in LA relative to MS underscores the tropical germplasm’s adaptation to warmer climates. Year-to-year variation within locations further emphasizes the dynamic nature of G × E interactions and the importance of multi-year data, ideally integrated with weather sensor information, for robust modeling and stable cultivar recommendations.

PhenoStream also enabled the entry-level analysis, providing a deeper layer of insight, moving from general species responses to specific genotypic performance. The case of *Miscanthus × giganteus* entry UI12-001-007, which consistently ranked high in AL and IL but low in MS, is a compelling example of a G×E interaction. This suggests that while this entry is broadly adapted to temperate conditions, its performance declines in the specific environment of MS, possibly due to particular environmental factors. Similarly, the intermediate performance of energycane entry Ho14-9213 in MS, despite the general superiority of the species in LA, points to genetic variability in environmental adaptation even within a tropical species. The use of pairwise distance and reaction norm analyses provided an initial step for visualizing these complex interactions, offering a method for breeders to identify genotypes with either specific adaptation or broad stability.

The AI-based inference components, though presented as GUI tools within the pipeline, represent a significant step forward. The use of a ViT for yield prediction from temporal image sequences is a novel application in crop phenotyping, moving beyond traditional machine learning and vegetation indices feature selection (Liang et al., 2025). This approach has the potential to capture complex spatiotemporal patterns related to biomass accumulation that are missed by conventional methods. Similarly, the application of a self-supervised SimCLR is scarcely reported (Shah et al., 2026) and novel for determining flowering dynamics. By learning feature representations without extensive labeled data first, this method can overcome the scarcity of annotated training images, a major hurdle in applying traditional (Zhang et al., 2021) or deep learning (Varela et al., 2022a) to new phenotyping tasks. This application enables mapping flowering dynamics and their progression across locations, offering valuable insights for breeders and agronomists seeking phenological understanding of entry adaptation to local environments, and thus provides a practical solution in this context.

Further downstream analysis on the environmental plasticity of bioenergy species and hybrids are consistent with previous studies that have documented G × E interactions in these crops (Shaik et al., 2026a), (Shaik et al., 2026b). However, previous work has often been limited to pairs of locations or only a few time points due to the logistical constraints of manual phenotyping. The value of our study lies in the high-resolution, multi-temporal, and multi-location dataset that provides a much richer and holistic picture of these dynamics. The modular integration and value of the PhenoStream was demonstrated by modeling G × E interaction in a case of study. When considering the multi-year interaction term as part of the model, yield prediction demonstrated superior performance than simplified models, hence helping to optimize the allocation of resources (Jarquin et al., 2020). In our case, the use of phenomics as proxy for predicting plant performance is justified by the complexity of phenotyping dry biomass yield.

This automated pipeline has immediate and practical implications. It enables the management of large-scale, distributed phenotyping trials with efficiency previously unavailable. Breeders can now make rapid, data-driven selections based on temporal growth patterns and environmental responses rather than single manual time-point measurements.

Future directions aim to address current limitations and enhance the functionality of the proposed cyberinfrastructure. First, although the selected UAV platform provides a reasonable cost-effective solution, certain limitations remain. Despite RTK capabilities, the aerial system exhibited limited repeatable accuracy for georectifying successive dates of flights, requiring the development of a custom post-processing solution. In particular, co-registration (i.e., geometrical alignment) but also vertical alignment (i.e., z dimension) of orthomosaics and CSMs between successive dates is not consistently solved, while global and local distortions typical limitations of non-RTK systems are. Second, expanding the integration of high-quality ground-truth data for traits such as chemical composition, disease, phenology and yield is essential for calibrating and validating remote sensing-based predictions. Consequently, algorithm refinement, particularly through domain adaptation techniques to improve performance across environments and species-depends on a robust database of imagery paired with corresponding ground-truth measurements, an effort that is currently underway. Finally, the development of ‘breeding-informing’ modules as part of the cyberinfrastructure that can directly translate spatiotemporal data into selection indices or genomic selection models will further reduce the gap between phenotyping and genetic improvement.

## CONCLUSION

We have developed and validated an integrated automation and cyberinfrastructure that effectively reduces data silos and minimizes latency in high-throughput phenotyping. By applying this pipeline to a practical use case as a regional network of field trials, we demonstrated its capacity to generate insights in the spatiotemporal dynamics and traits inference of bioenergy crops, successfully differentiating the effects of genotype, environment, and their interaction. The move from generalized species responses to entry-specific performance profiles underscores the power of this approach for precise decision-making. This work provides a scalable foundational work for the next generation of data-driven phenotypic pipelines, directly addressing the need to accelerate the development of high-yielding and adapted cultivars cost-effectively.

## ACKNOWLEDGMENTS

We thank the operational staff at each of the research stations for their support in field management, logistics, training, and data collection activities. We also thank, professor Brian Baldwin for his support as part of the Mississippi State University team.

## Author contributions

S.V. conceived the study and wrote the manuscript. A.D.B.L. provided critical feedback on the manuscript structure. S.V. and J.R. implemented the computational infrastructure. E.S., X.Z., A.H., J.M., E.C., X.K., and B.L. established and maintained the field trials, collected ground-truth data, and provided feedback on the manuscript. J.R., D.A., Y.Z., C.L., J.M., and B.L. conducted UAV-based aerial data collection. Y.Z. contributed to specific coding components of the data stream. D.J., S.D.P., and S.K. implemented the downstream plant breeding analyses and provided feedback on the manuscript.

## DATA AVAILABILITY

The datasets and code used during the current study are available from the corresponding authors upon request.

